# Introducing conflict resolution and negotiation training into a biomedical sciences graduate curriculum

**DOI:** 10.1101/2021.04.05.438459

**Authors:** Michael D. Schaller, Amanda Gatesman-Ammer

## Abstract

Analysis of the biomedical workforce and graduate education have produced recommendations for modifications of pre-doctoral training to broadly prepare trainees for wider ranging scientific careers. Increased exposure to career opportunities and development of training in professional skills is widely recommended, but details of implementation are not widely available. In alignment with these recommendations, we have incorporated professional skills training into the biomedical science graduate curriculum at West Virginia University. An important component of the training is developing conflict resolution and negotiation skills. This training will provide useful skills for academic careers, non-academic careers and life situations outside of the workplace. Conflict resolution/negotiation skills are also relevant in managing issues in diversity, equity and inclusivity. We report our experience in developing this component of the training program, provide an overview of the approach to delivery and practice of skills, and provide an analysis of the reception and effectiveness of the training.

## Introduction

Our paradigm of graduate education for biomedical scientists was designed to train doctoral students to perform research and prepare them for an academic career. The number of students graduating with a Ph.D. each year has steadily increased since 1990, while the number of tenured/tenure-track faculty has remained constant (Ghaffarzadegan, Hawley, Larson, & Xue, 2015; National Academies of Sciences Engineering and Medicine, 2018a, 2018b; National Institutes of Health, 2012). Consequently, new Ph.D.s are entering the workforce in non-tenure track academic positions, research positions in industry or government, teaching intensive and administrative positions, and other non-research-intensive positions (National Institutes of Health, 2012; Sinche et al., 2017). Further, surveys of doctoral students suggest that a large percentage of trainees are interested in pursuing non-academic careers/non-research careers (Fuhrmann, Halme, O’Sullivan, & Lindstaedt, 2011; Roach & Sauermann, 2017; Sauermann & Roach, 2012; St Clair et al., 2017). Reports from the National Institutes of Health and the National Academies of Science, Engineering, and Medicine recommend modification of graduate student training to prepare them for a broader range of careers (National Academies of Sciences Engineering and Medicine, 2018a, 2018b; National Institutes of Health, 2012). These recommendations include increasing transparency of outcomes of graduate programs, increasing opportunities to explore different careers, and providing training in professional skills. Interestingly, a large survey of Ph.D. recipients indicates that similar soft/transferable skills are required by all Ph.D.s, regardless of their traditional or non-traditional career path, indicating that targeted training in these areas are beneficial for all Ph.D. candidates (Sinche et al., 2017). The NIH has broadly invested in this area via the BEST (Broadening Experiences in Scientific Training) program, which supported 5-year programs at 17 institutions to establish and disseminate best practices in career development and professional development (Meyers et al., 2016). The NIGMS has also introduced guidelines for modifications to training programs to incorporate career development and professional development activities (Singh, Gammie, & Lorsch, 2016).

Recommendations of desired professional skills competencies for doctorates have emerged from student surveys, employer surveys, committee reports, and workshops (Denecke, Feaster, & Stone, 2017; Fuhrmann et al., 2011; Hitchcock et al., 2017; National Academies of Sciences Engineering and Medicine, 2018a; Sinche et al., 2017; Verderame, Freedman, Kozlowski, & McCormack, 2018). A compilation of these recommendations is reported in Table 1. One competency is designed to address the NIH and NASEM recommendation to increase the opportunity for trainees to explore career opportunities (i.e. career planning/awareness). Many of the BEST programs have successfully created mechanisms to address career awareness and planning (Infante Lara, Chalkley, & Daniel, 2020). Some competencies obviously translate to important skills in a research-intensive scientific career, including communication, project management, and critical thinking, while others are comprised of a complex set of skills that are less obviously linked, such as leadership, managing others, and teamwork. A common thread to the latter competencies is personnel management and interpersonal relationships. Building relationships, managing difficult conversations, and dealing with conflict are specific skill sets underlying these competencies. In response to these recommendations, we have incorporated professional skills development into our first-year biomedical sciences graduate curriculum (as part of a larger course), emphasizing these specific skill sets. A major emphasis in these sessions is conflict resolution and negotiation, skills that transcend these competencies. Herein, we describe our experiences in developing these sessions and offer a template for similar instruction.

**Table 1.** List of Professional Skills Competencies Compiled from the Literature.

| Competency | References |
| --- | --- |
| Leadership | (Denecke et al., 2017; Fuhrmann et al., 2011; Hitchcock et al., 2017; Verderame et al., 2018) |
| Communication | (Denecke et al., 2017; Fuhrmann et al., 2011; Hitchcock et al., 2017; National Academies of Sciences Engineering and Medicine, 2018a; Verderame et al., 2018) |
| Project management | (Denecke et al., 2017; Fuhrmann et al., 2011; Hitchcock et al., 2017; National Academies of Sciences Engineering and Medicine, 2018a) |
| Teamwork | (Denecke et al., 2017; Hitchcock et al., 2017; National Academies of Sciences Engineering and Medicine, 2018a; Sinche et al., 2017; Verderame et al., 2018) |
| Critical thinking | (Denecke et al., 2017; Hitchcock et al., 2017; Verderame et al., 2018) |
| Collaboration | (Denecke et al., 2017; Hitchcock et al., 2017; National Academies of Sciences Engineering and Medicine, 2018a; Sinche et al., 2017) |
| Time management | (Denecke et al., 2017; Sinche et al., 2017) |
| Setting visions and goals | (Sinche et al., 2017) |
| Managing others | (Sinche et al., 2017; Verderame et al., 2018) |
| Career planning/awareness | (Sinche et al., 2017) |
| Networking | (Fuhrmann et al., 2011) |
| Interculture competency | (Denecke et al., 2017) |
| Problem solving | (Denecke et al., 2017) |
| Resilience | (Denecke et al., 2017) |
| Entrepreneurship | (Denecke et al., 2017) |
| Science policy | (Denecke et al., 2017) |

## Materials and methods

The West Virginia University IRB approved the study. This study was partially retrospective and partially prospective. IRB approval numbers are WVU Protocol#: 2001865972 and WVU Protocol#: 2004977501. Informed consent was in writing and all information was anonymous.

Evaluation of effectiveness of training was similar to recommendations by Denecke et al. (Denecke et al., 2017), which align with the principals of the Kirkpatrick Four Level Model of Evaluation (Kirkpatrick, 2006). While originally developed as a tool for evaluation of human resources training, this evaluation model is widely used and has been adapted to assessment of higher education (Praslova, 2010). Upon completion of training, students completed a questionnaire about their perceptions of training (Level 1 evaluation – reaction) (Table 2). This was constructed as a formative assessment of the training sessions. At the end of the semester, students were asked how they would respond to different scenarios requiring conflict resolution/negotiation skills to measure what they had learned (Level 2 evaluation – learning) (Table 3). Several months later, students were surveyed to determine if they used some of these skills and/or witnessed situations where these skills would be useful, i.e. if their behavior had changed (Level 3 evaluation – behavior) (Table 4). We have not attempted to evaluate the fourth level (results), which would require direct measurements of the success of trainees at managing conflict and negotiating. The timing of training sessions and surveys for workshops and classes is illustrated in Fig 1. Responses to all of these surveys were anonymous. Therefore all of the analyses compared responses between individuals.

**Table 2.** Survey Questions for Student Reaction to Training (Level 1).

|  |
| --- |
| **Students were asked to score on Likert scale (1 = strongly disagree, 5 = strongly agree)** |
| **Q1.** The sessions provided me with new information about conflict resolution |
| **Q2.** The role-playing activities helped me prepare for conflict resolution. |
| **Q3.** This course has provided me the opportunity to reflect on my personal approach to conflict. |
| **Q4.** This course has provided me insight into how others might approach conflict. |
| **Q5.** This course has provided me with strategies that I will try to use to resolve conflict at work/home. |
| **Q6.** The sessions provided me with new information about negotiation. |
| **Q7.** The role-playing activities helped me prepare for negotiations. |
| **Q8.** This course has provided me with strategies that I will try to use when I negotiate in future. |

**Table 3.** Questions to Evaluate Student Learning from Training (Level 2)

|  |
| --- |
| **Q1.** When you enter into negotiation, what can you do to strengthen your position/increase your leverage? |
| **Q2.** You are about to negotiate and you would like to use principled negotiation. What does this mean to you? |
| **Q3.** Make a statement/ask a question illustrating active listening. |
| **Q4.** You and a colleague are working collaboratively on a side project and the balance of the work is not being equally shared. You have done most of the planning and execution of the experiments thus far. This project will be productive if your colleague fully engages. In a few sentences, begin the conversation about balancing the work with your colleague. |
| **Q5.** Your colleague says, “Last night, you left the equipment I needed this morning in the sink and dirty again. You never clean up after yourself. Why do you insist on being such a pig? ”. How do you respond to reframe the attack? |
| **Q6.** In a difficult conversation, your colleague goes silent. What do you need to do and how can you do it? |
| **Q7.** In a difficult conversation, your colleague becomes very sarcastic and insulting. What would you say to “surface/name” the attack and reframe the conversation? |
| **Q8.** A student working under your supervision demonstrates great technical skills, but keeps very poor notes. What would you say to the student to encourage him/her to keep better notes? |
| **Q9.** What is one strategy you can use when you are negotiating against power? |
| **Q10.** How would you respond when your opponent is stonewalling in a negotiation? |

**Table 4.** Survey Statements/Questions to Explore Student Behavior (Level 3)

|  |
| --- |
| 1. I have used concepts from conflict resolution/negotiation in my professional and/or personal interactions with others. (yes/no). |
| 2. What concepts from these sessions did you use? |
| 3. I have observed situations where the use of concepts from conflict resolution/negotiation would have been beneficial. (yes/no). |
| 4. What concepts from these sessions do you think would have been beneficial in that/those situation(s)? |
| 5. In the future, I intend to use skills from the conflict resolution/negotiation sessions in my interactions with others. (yes/no). |
| 6. What concepts do you think will be useful to you in future? |

**Fig 1.**
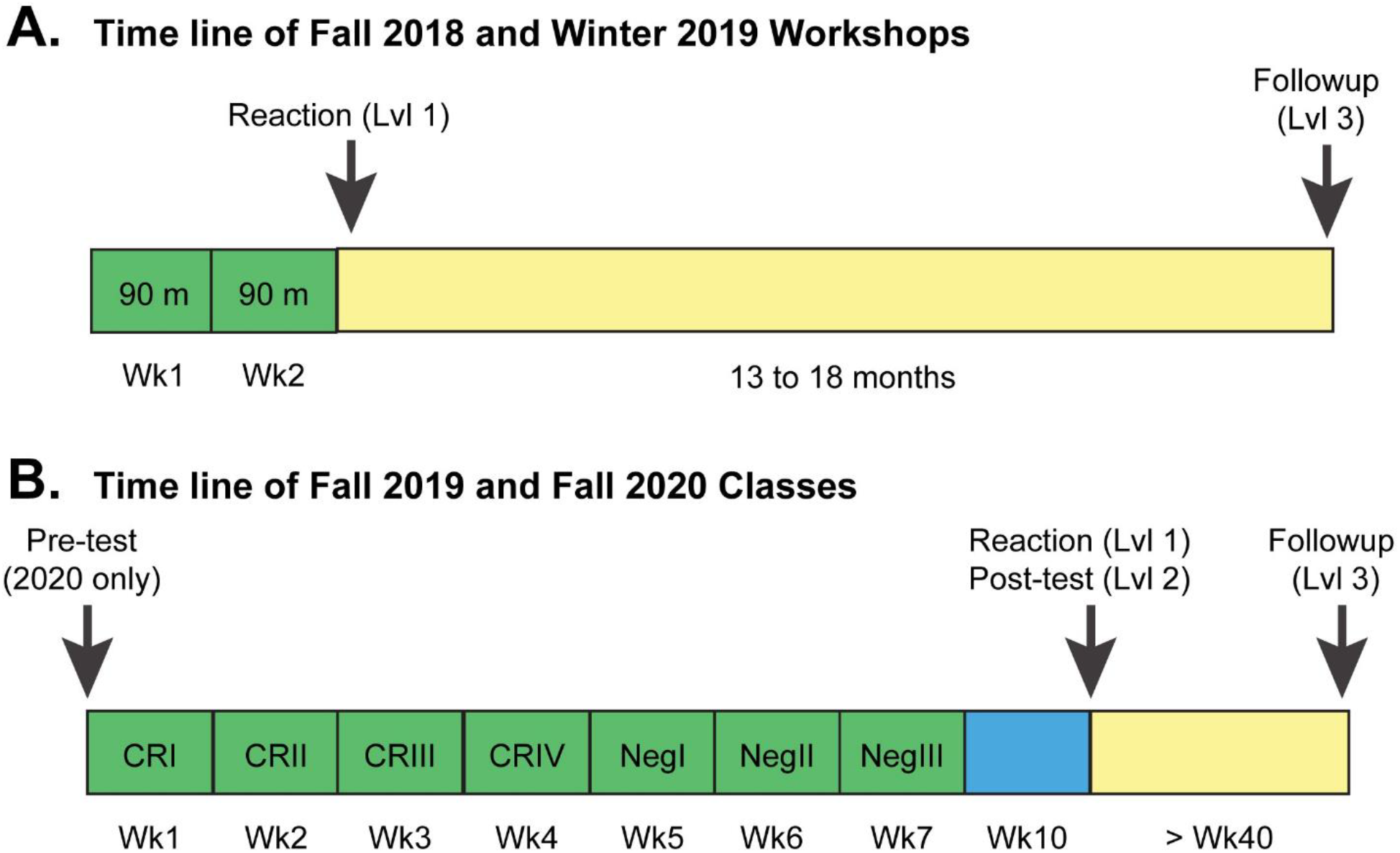
Number of Trainees and Respondents to Surveys. Two formats, a workshop and classroom training, were used to provide training in conflict resolution. The timing of training and surveys for workshops (A) and classroom training (B) is illustrated. Training sessions are indicated in green and intervening times until administration of surveys in blue and yellow. The timelines are not drawn to scale.

The ten questions to evaluate learning in post-tests in 2019 and 2020, and to evaluate baseline knowledge in a pre-test in 2020, were independently scored by both instructors using a common rubric. Scores were averaged for statistical analysis. Brown-Forsythe and Bartlett’s tests revealed homogeneity of variance between the samples. However, Shapiro-Wilks, Anderson-Darling, D’Agostina & Pearson and Kolmogorov-Smirnov tests for normal distribution revealed that these scores were not Gaussian and therefore a non-parametric statistic was used. The Kruskal Wallis H test and Conover and Dunn posthoc tests were used to evaluate differences between scores. (The Kruskal Wallis H test is the non-parametric equivalent of an ANOVA analysis.)

## Results

### Exploring instructional platforms

The impetus to develop training in conflict resolution came from participation in the Train the Trainer Workshop sponsored by the Office of Intramural Training at the National Institutes of Health. Training in conflict resolution at WVU was developed and performed using two different formats: initially as a workshop and later as a classroom component of the formal first year curriculum. The number of attendees and respondents to surveys for workshops and classroom training are indicated in Table 5. Workshops were held over the course of two weeks as two ninety-minute sessions. The first workshop (October 2018) targeted trainees affiliated with T32 training programs and contained a didactic session followed by a role-playing session the following week. The second workshop (February 2019) targeted students in two biomedical science graduate programs (the Biochemistry and Molecular Biology and Clinical and Translational Sciences Graduate Programs) and consisted of two mixed sessions containing didactic and role-playing components. The role-playing exercises utilized real situations of faculty-student conflict, solicited from faculty and students at WVU. While the workshops were well received, student commitment to participation flagged as the number of participants was significantly less than the number of students registered to participate. Anecdotal evidence from discussion with students who participated in the workshops suggested that training in conflict resolution would be useful to all graduate students and that training early in the curriculum would be most beneficial. Conflict resolution was incorporated into a professional skills development component for the first-year graduate curriculum for fall 2019.

**Table 5.** Number of Attendees and Respondents to Surveys.

| Format | Registered | Attendees | Level 1 Response | Level 2 Response | Level 3 Response |
| --- | --- | --- | --- | --- | --- |
| Workshop 2018 | 15 | 10 | 5 | na | 9<sup>a</sup> |
| Workshop 2019 | 15 | 10 | 5 | na |  |
| Fall 2019 Class | 28 | 28 | 28 | 28 | 19 |
| Fall 2020 Class | 30 | 30 | 26 | 26 | 19 |

<sup>a</sup>Level 3 follow up survey did not distinguish which workshop respondents attended.

The professional skills development component of the first-year curriculum was designed to include didactic/interactive sessions on communication, lay communication, resilience, networking, working in teams/collaboration, conflict resolution, and negotiation (Table 6). Interestingly, six of the nine topics covered in our Professional Skills session were also incorporated in the URBEST Program’s Leadership and Management for Scientists at the University of Rochester (Baas, Dewhurst, & Peyre, 2020). Each session was 50 minutes in duration. We also offer a 1-credit course with identical content that runs concurrently for students not enrolled in this curriculum, including more senior students and students from other departmental programs. Through this mechanism, a broader impact upon the graduate programs at WVU is anticipated. Conflict resolution was a central theme of these sessions, since these skills crossed a number of recommended professional skills competencies, including leadership, project management, teamwork, collaboration, and managing others. Negotiation, which shares many concepts with conflict resolution, was also incorporated. Our philosophy is to use collaboration to resolve conflict constructively and to use principled negotiation strategies. Each session was a mixture of didactic presentation of concepts interspersed with interactive/role-playing activities. Distribution of the sessions over seven weeks was intentional to reinforce concepts and engage in role-playing over an extended period of time to promote changes in the participants’ behavior in conflicting situations. A conflict resolution workshop at Michigan State University is not part of the formal curriculum, but is similarly extended over six sessions (J. L. Brockman, Nunez, & Basu, 2010). This strategy is based on the concepts of retrieval learning (role playing), spacing out practice (over multiple sessions), and varying practice (interleaving conflict resolution and negotiation) as effective methods of learning (Brown, Roediger III, & McDaniel, 2014). The Fall 2020 iteration of the course utilized a virtual format, due to the COVID-19 pandemic.

**Table 6.** Professional Skills Development Topics Incorporated.

| Session | Topic |
| --- | --- |
| **Communication Skills** |  |
| 1 | Prof Dev I - Communication - How to give a talk |
| 2 | Prof Dev II - Communication - to non-experts |
| **Personnel Management/Interpersonal Interactions** |  |
| 3 | Prof Dev III - Conflict Resolution - theory |
| 4 | Prof Dev IV - Conflict Resolution - practice |
| 5 | Prof Dev V - Conflict Resolution 3 |
| 6 | Prof Dev VI - Crucial Conversations |
| 7 | Prof Dev VII - Negotiations - theory |
| 8 | Prof Dev VIII - Negotiations - practice |
| 9 | Prof Dev IX - Shadow Negotiations |
| **Beneficial Skills** |  |
| 10 | Prof Dev X - Networking |
| 11 | Prof Dev XI - Perseverance/Resilience |
| **Team and Management** |  |
| 12 | Prof Dev XII - Entrepreneurship |
| 13 | Prof Dev XIII - Working in Teams |
| 14 | Prof Dev XIV - Collaboration |
| 15 | Prof Dev XV - Project management |

### Organization of conflict resolution/negotiation sessions

The conflict resolution/negotiation sessions were comprised of four components interspersed throughout the session. The first was didactic, where concepts, strategies, and tactics for managing conflict/negotiation were presented to the students. The second was interactive, where the students contributed to in-class discussion using on-line polling tools. The third was an illustration of a relevant situation acted out by the instructors. These were first done to demonstrate the ‘wrong’ way to manage the situation and how this approach could spiral out of control. The situation was acted out a second time demonstrating a specific strategy or tactic to manage conflict in that scenario. The fourth component was role-playing by the students. The scenarios for role-playing were solicited from students and faculty at the Health Sciences Center at WVU and were drawn from their actual experiences. These activities provided an opportunity to practice specific conflict resolution skills and the student playing the ‘opponent’ in the exercise was given specific instructions to resist. The role-playing activities were considered important to begin training the students to modify their behavior to improve their ability to resolve conflict. Active learning activities like these are frequently utilized in professional education workshops providing training in conflict resolution (Klomparens, Beck, Brockman, & Nunez, 2008; Shrader & Zaudke, 2015; Welch, Jimenez, & Allen, 2013; Wolfe, Hoang, & Denniston, 2018).

Each of the sessions was designed to deliver a few lessons related to conflict resolution/negotiation and some skills for students to learn to manage conflict (Table 7). The lessons and skills were developed from a number of sources (Table 7). The first sessions focused upon basic skills and subsequent sessions built upon these skills to elaborate on more complex scenarios, strategies, and tactics. When transitioning to the negotiation sessions, twenty-four concepts from conflict resolution were briefly reviewed, as these were also essential concepts for successful negotiation (Table 8). The interactive exercises with the class were interspersed with the didactic parts of the lectures. One example is an activity used in a session about emotions in conflict, where strategies to reframe emotions and defuse the opponent’s emotions are discussed. In the exercise, “scenes from a hat”, one student draws an inflammatory statement (authentic statements heard by and solicited from graduate students) from a hat and reads the statement to another student. For example, “Why don’t you ever help out with laboratory grunt work? You always use up all the reagents and you never make them! I am so sick of this!”. The other student responds quickly to reframe this personal challenge to focus on the underlying issues, rather than on the persons involved. For example, “You are right that there is an issue of keeping common reagents stocked. Perhaps we could discuss ways to work as a team to restock.” A second example of using interactive activities to develop concepts come from the first session of negotiation, which covers the topics of preparing for negotiations, thinking outside the box and developing a strong BATNA (best alternative to a negotiated agreement). At the beginning of the session, the students are asked to assume the role of an Assistant Professor whose boss has just asked them to take on the role of Director of Graduate Studies. The students are polled anonymously for their opinions of the pros and cons of taking the position. Responses were projected in real time, providing the students insight into the thoughts of their colleagues and providing collective lists. The session then moved into a didactic component discussing preparation and performing research prior to the beginning of a negotiation. The students were then anonymously polled for factual information they would like to collect. The session returned to a didactic discussion of interests and inventing options. Finally, the students were polled for concessions they could ask for in return for taking on this assignment. Most of the responses from the students were thoughtful and reasonable, suggesting they took the opportunity to seriously work through the exercise.

**Table 7.** Details of Conflict Resolution/Negotiation Sessions.

| Session | Lessons | Skills | References |
| --- | --- | --- | --- |
| Conflict Resolution 1 | • Constructive conflict<br>• Get to the real problem<br>• Separate issues<br>• My truth ≠ your truth | • Active listening<br>• Situation/action/impact feedback | (Cloke & Goldsmith, 2011; Deutsch, 2014; Stone, Heen, & Patton, 2010) |
| Conflict Resolution 2 | • Conflict can be emotional<br>• Stories generate emotions | • Reframing your emotions<br>• Defusing your opponent's emotions | (Cloke & Goldsmith, 2011; Deutsch, 2014; Patterson, Grenny, McMillan, & Switzler, 2012; Sandy, |
|  |  |  | Boardman, & Deutsch, 2014; Stone et al., 2010) |
| Conflict Resolution 3 | • Managing feelings and identity<br>• Focus on contribution, not blame<br>• Intentions – don’t assume, careful of impact | • Using 3<sup>rd</sup> story to start<br>• Inquiring, paraphrasing, acknowledging<br>• “Me-me, and” | (Stone et al., 2010) |
| Conflict Resolution 4 | • Keeping and regaining focus<br>• “silence” & “violence” impede dialogue – need to make it safe<br>• Restoring safety – mutual purpose & mutual respect | • Contrasting<br>• STATE your path – share facts, tell story, ask their story, talk tentatively, encourage testing<br>• Ask/mirror/paraphrase/prime | (Patterson et al., 2012) |
| Negotiation 1 | • Principled negotiation<br>• Preparation – facts and interests<br>• BATNA | • Inventing options – thinking outside the box<br>• Improving your BATNA | (Fisher, Ury, & Patton, 2011; Kolb & Williams, 2006; Siedel, 2014; Ury, 2007) |
| Negotiation 2 | • Anchoring<br>• Cognitive dissonance<br>• Exploiting differences | • Expressing yourself<br>• Asking questions<br>• Maintaining engagement | (Fisher et al., 2011; Kolb & Williams, 2006; Shell & Moussa, 2007; Siedel, 2014; Ury, 2007) |
| Negotiation 3 | • Shadow negotiations<br>• Resistance to tricky tactics | • Redirecting or turning attacks<br>• Reframing | (Fisher et al., 2011; Kolb & Williams, 2006; Ury, 2007) |

**Table 8.** Twenty-four Conflict Resolution Strategies used in Negotiation.

|  |  |  |
| --- | --- | --- |
| Separate people from problems | Use active listening | Surface the attack |
| Focus on interests not positions | Speak to be understood | Know your hot buttons |
| Empathy | Speak about how you feel | Agree where you can |
| Don't blame | Build a relationship | Show that you heard them |
| Don't assume intentions | Ask open ended questions | Acknowledge |
| Understand your emotions | Focus on future, not past | Reframe |
| Prepare for challenges to identity | Be hard on the problem, soft on the people | Resist attack/counterattack spiral |
| Apologize when appropriate | Don't say "but", say "and" | My truth is not your truth |

The role-playing exercises were very important components of these sessions. The demonstrations by the instructors were very well received by the students. These were semi-scripted to ensure that the exercises properly illustrated the points intended. One effective demonstration addressed managing your emotions and regaining control under difficult circumstances. The scenario demonstrated the interaction between a critical faculty member and a student making a public presentation. As the faculty member continues to criticize, the student responds emotionally, either withdrawing or lashing out at the faculty member. The demonstration continues after the student lashes out to illustrate how the student can regain control by apologizing for the outburst and turning the attack of the faculty member, e.g. “From your comments, I can see that there may be additional considerations that you could help me with. Perhaps we could meet and discuss these issues at a later date rather than in the middle of my presentation.” A second example of instructor demonstrations illustrated the reason for asking open ended questions during negotiation. The demonstration begins, “Can we change this policy?” The response, “No.” At this point, the negotiation is effectively over and this exercise exemplifies that negotiations can end quickly if questions are asked in the wrong fashion. The follow up demonstration uses open ended questions. “What is the purpose of this policy?”, “Who created this policy?”, “Why not modify the policy to take into account this situation?”. Illustrating different examples by demonstration provides an alternative to didactic presentation of the concepts. Perhaps the most important and best received exercises were the role-playing activities for students, since this provided them the opportunity to practice different approaches in conflict resolution and negotiation. These activities were designed to practice the specific skill sets covered in class and the ‘opponent’ in the discussion received instructions to resist and raise objections to extend the opportunity to practice different skills. Many scenarios were developed for the workshops and the class. The scenarios utilized for conflict resolution exercises are shown in the supplemental material. During these exercises, the instructors circulated among the students to observe their interactions, identify good examples to point out to the class, and provide suggestions if the students were struggling. Immediately following the exercise, the class and instructors discussed statements and approaches from the exercise that appeared impactful.

### Incorporation of microaggression

The conflict resolution sessions contain strategies for engaging in difficult conversations and some of these concepts are very relevant to managing difficult conversations around diversity and inclusion. The negotiation sessions contain strategies to deal with tricky or challenging bargaining tactics and include strategies to counter demeaning, exclusionary, undermining and harassing behavior during negotiations. In the Fall 2020 version of the course, a conflict resolution discussion on microaggression was implemented. Public service announcements from MTV and YouTube videos were used to illustrate examples of microaggression that were discussed as part of the class. The frequent benign intent of microaggressive statements was contrasted with the detrimental impact upon the recipient of the comment. Strategies to respond to microaggressions, by both the recipient and a bystander were presented. Students were asked to reflect on microaggression and write about a fictional incidence of microaggression explaining how the recipient might respond and how a bystander might respond to the microaggression.

### Effectiveness of Training

#### Student reception of the workshop/course

Upon completion of the workshops, students were requested to take an online, anonymous survey. Approximately 3 weeks after completion of the conflict resolution/negotiation sessions in the first-year curriculum, the students were asked to complete an online, anonymous survey to measure their reception of the sessions. The workshop survey contained five statements related to the objectives and activities of the workshop, and the professional skills development course included three additional statements related to the negotiation sessions (Table 2). Participants were asked if they agreed with the statements using the Likert scale to indicate strength of agreement (5 = strongly agree) or disagreement (1 = strongly disagree). The results are shown in Fig 2. The students were largely in agreement that the workshop/course provided new information, helped them prepare for conflict and negotiation, provided insight into their approach and others approaches to conflict, and provided useful strategies for use in the future. The average scores were remarkably uniform across the different iterations of these sessions. The reaction of students to these sessions suggest that they provide new information about conflict resolution/negotiation and some preparation for managing conflict and negotiating in the future. An interesting observation is the lowest Likert scores on these surveys related to the usefulness of the role-playing activities in preparation for conflict resolution and negotiation.

**Fig 2.**
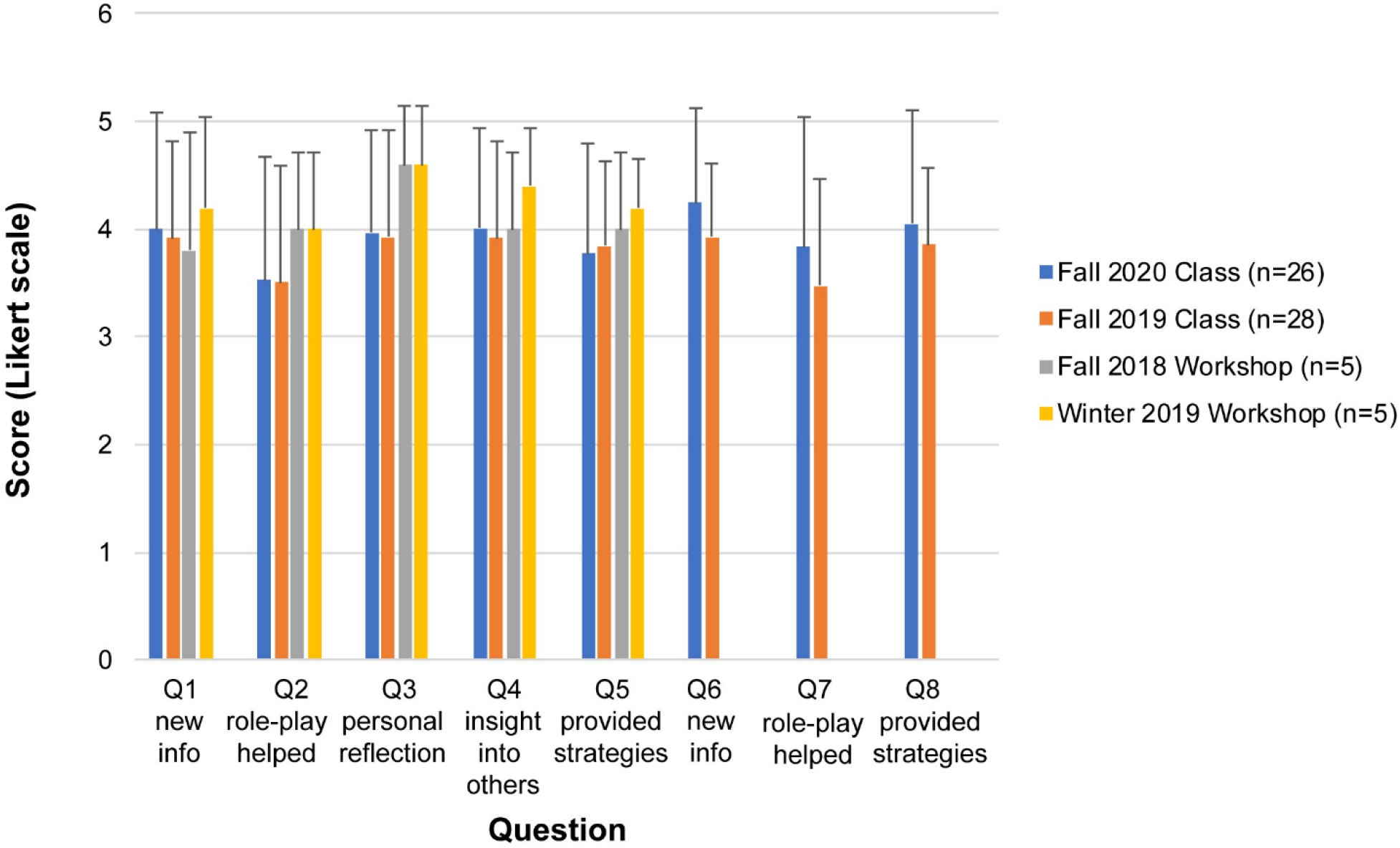
Student Responses to Workshop/Class Survey. Students were asked to respond to questions on the survey (see Table 2) on the Likert scale. Questions Q1 to Q5 were related to conflict resolution and questions Q6 to Q8 were related to negotiation. The average score +/-standard deviation is plotted. The average score for each iteration of the conflict resolution/negotiation training sessions is shown. n = number of respondents

The students were also asked to describe the best part(s) of the workshop/professional skills sessions (Table 9). Given the numerical response to the statement about the usefulness of role-playing (Fig 2), it was surprising that the students overwhelmingly indicated that this was the best part of the course. Distant seconds in the students’ responses including watching simulated arguments/negotiations, the didactic content and learning about personal approaches to conflict.

**Table 9.**
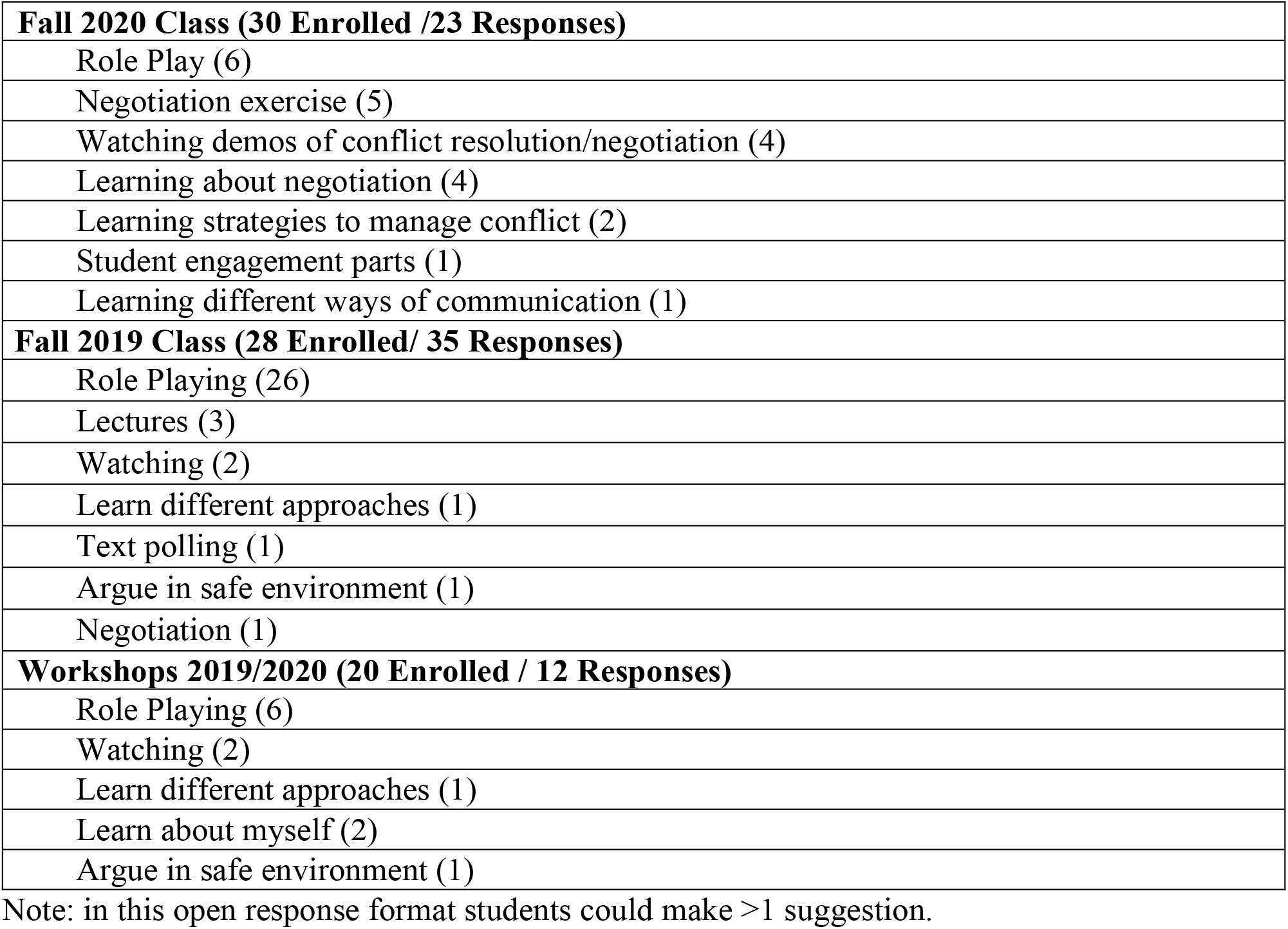
Best Parts of the Conflict Resolution/Negotiation Sessions.

Students were also asked to make suggestions for improvement for the sessions, although there was little consensus (Table 10). A group of students suggested incorporating more role playing, while another group of students suggesting reducing the amount of role playing. Several students suggested incorporating more realistic scenarios, although the scenarios used came from a solicitation to students and faculty to provide examples of conflicts in which they were involved or had observed. The groups/partners for these sessions in fall 2019 were not assigned, but rather self-assembled, and one suggestion for improvement was to change up the groups to provide new partners for role-play. Students were randomly assigned for role-playing in the virtual classroom in fall 2020. One suggestion was to increase the number of facilitators for the role-playing sessions. The primary method of delivery of content was didactic and the use of additional media(e.g. videos to illustrate points) was suggested. One suggestion emerging from a workshop was the incorporation of situations of co-worker conflicts in the role-playing sessions. This was done in the classroom sessions on conflict resolution and these appeared to be well received.

**Table 10.** Suggestions for Improvement for the Conflict Resolution/Negotiation Sessions.

|  |
| --- |
| **Fall 2020 Class (30 Enrolled/ 15 Responses)** |
| Less role playing (2) |
| Don't force people to talk (2) |
| Increased engagement in some sessions (1) |
| More on how to manage aggressive opponent (1) |
| Provide more information in advance (1) |
| More negotiation practice (1) |
| More role playing (1) |
| Indicate extra sessions on syllabus (1) |
| More backstory for negotiation exercise (1) |
| Negotiation should be later in grad school (1) |
| Different, more relevant negotiation exercise (1) |
| More activities, less didactic (1) |
| **Fall 2019 Class (28 Enrolled/ 23 Responses)** |
| More realistic scenarios (3) |
| More role-playing (7) |
| Less role-playing (4) |
| Reduce number of sessions (1) |
| Use multimedia, including videos (1) |
| Change up groups (1) |
| Expertise in negotiations (1) |
| More time negotiation strategies (1) |
| More facilitators to monitor (1) |
| Reduce didactic portion (1) |
| Reduce conflict sessions (1) |
| Improve structure negotiation exercise (1) |
| **Workshops 2019/2020 (20 Enrolled / 9 Responses)** |
| More role playing (2) |
| Distribute materials (1) |
| Demo conflict with different personalities (1) |
| More professors (1) |
| Principle investigator involvement would provide insight to participants (1) |
| Conflict with co-workers (1) |
| More demonstration conflicts (1) |
| Multi-dialogue, multi-strategy (1) |

Note: in this open response format students could make >1 suggestion.

#### Student learning in the course

As part of the survey to evaluate student reception of the course, the students were also asked ten questions to determine if they retained specific concepts on conflict resolution and negotiation (Table 3). This was not a formal test, was administered anonymously and the students were instructed that there was no need to study in advance. The intent was to measure the quality of information that they retained from the training sessions and not their ability to master the subject for an exam. Both instructors used a common rubric to grade each student’s responses. The scores of the two instructors were averaged.

After completion of training in fall 2019, the students exhibited knowledge of strategies to manage conflict of interest and negotiation (mean score = 7.3 +/-2.1, median = 8) (Fig 3). After completion of training in fall 2020, the students scored lower on the assessment than the preceding cohort (mean score = 4.9 +/-2.1, median = 5). In fall 2020, a pre-test, which was identical to the post-test, was incorporated (mean score = 2.3 +/-1.6, median = 2)(Fig 3). The distribution of scores on each exam was not Gaussian and a non-parametrical statistical test, the Kruskal Wallis H test, was used to evaluate differences between the scores. The H statistic was 85.97, p<0.0001. Conover and Dunn posthoc tests indicated that the scores between all three tests were statistically different. The difference between the pre-test and post-test scores in 2020 suggests that the students learned concepts of conflict resolution and negotiation. It is uncertain if the difference in post-test scores between cohorts are due to different levels of knowledge prior to training (there was no pre-test in fall 2019), variability in learning between the cohorts or if it reflects the difference in learning in person and in a virtual format. As the goal of the training was to prepare students to better manage conflict and negotiation, overall performance in these evaluations is encouraging and suggests that these training sessions may have been effective.

**Fig 3.**
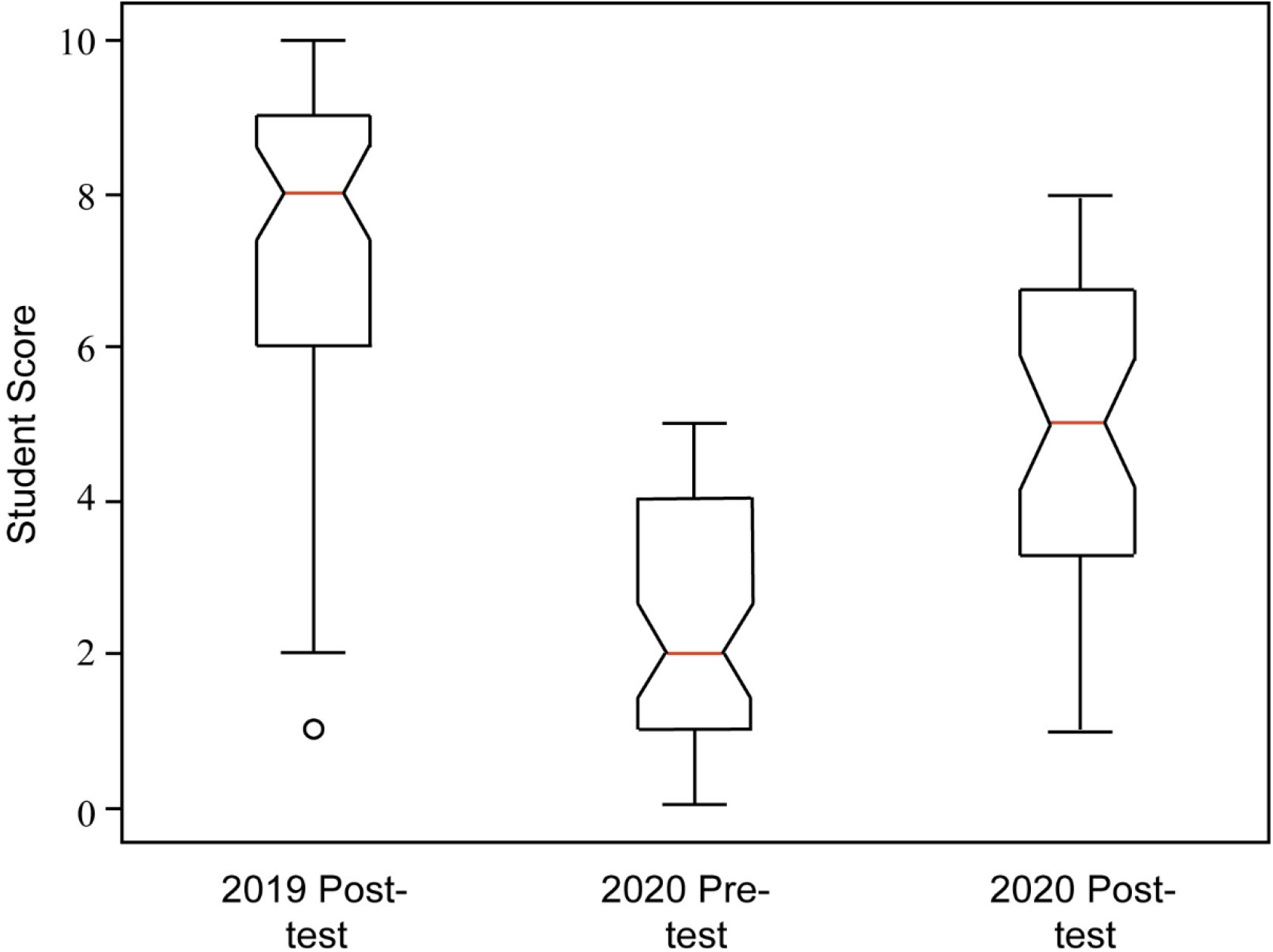
Student Scores on Pre- and Post-Tests. The students were asked 10 questions (see Table 3) to evaluate their knowledge of concepts and skills presented in the training sessions. A post-test was administered after the Fall 2019 and Fall 2020 iterations of the sessions and a pre-test was administered before the Fall 2020 sessions. The tests were scored independently by the two instructors using a common rubric. The test scores are presented as a box and whisker plot. The outlier in the Post 2019 test is indicated by a circle. The distribution of scores for each exam was non-Gaussian. The results were analyzed using the Kruskal Wallis H test (H = 85.97, p < 0.0001). Conover and Dunn posthoc tests were performed and indicated that the scores of each of the three tests were different.

#### Impact on the students – after the fact

Students were surveyed in March 2020 to determine if training in conflict resolution/negotiation had altered their behavior or attitude towards conflict resolution. This survey was 18 months after the first workshop, 13 months after the second workshop, and 5 months after the conflict resolution/negotiation sessions in the professional skills development class. All registrants/participants in the workshops who were still registered at WVU and all students who had taken the professional skills development class were surveyed. The survey consisted of three questions (Table 4), with a follow up to each to elaborate on the concepts from class that were relevant to the situation if the respondents answered in the affirmative. Students matriculating in Fall 2020 were surveyed in March 2021. The overall response rate for the surveys was 57.5%.

The majority of students who took a workshop had used concepts from the workshop in resolving conflicts and all of the students had observed scenarios where they felt that concepts from the course could have been applied (Fig 4). All of the students indicated an intent to utilize conflict resolution concepts in the future. The students who took the professional skills development sessions as part of the first-year curriculum were more junior than those who took the workshops, and there was less elapsed time between the conflict resolution sessions and the survey. The majority of these students had used concepts from the course and had observed situations where concepts could have been used (Fig 3 B and C). Most of the respondents stated their intention to use some of the concepts in future conflicts. While other influences upon students’ management of conflict cannot be excluded, these results are consistent with modified student behavior in response to training in conflict resolution.

**Fig 4.**
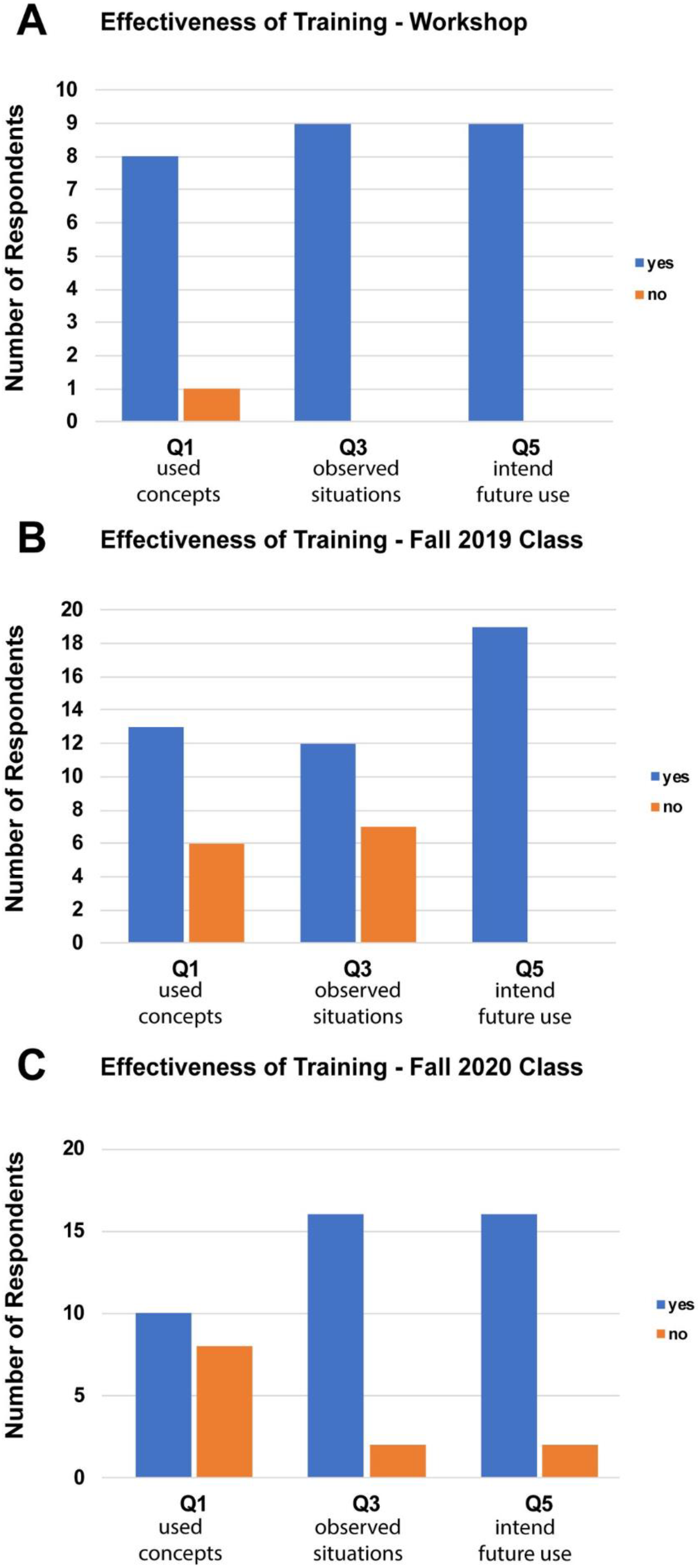
Student Responses to Follow up Surveys. Five to eighteen months after training, the students were surveyed to determine if they had applied concepts/skills from the training sessions or recognized situations where they might be employed. The surveys asked six questions (Table 4), three main questions and three follow up questions about the details of the concepts/skills used. The number of respondents answering ‘yes’ and ‘no’ to the three main questions (Q1, Q3 and Q5) are plotted for the students taking the training sessions in the workshop format (**A**) and the classroom format (**B** and **C**).

#### The other side of the equation – faculty training

In addition to providing conflict resolution training to graduate students, we have also conducted workshops on conflict resolution for faculty. These were single 60-to-90-minute sessions introducing some of the key concepts in conflict resolution and included interactive sessions designed to meet the needs of the group. In a mentor training workshop for faculty preceptors of doctoral students the interactive exercises focused upon conflict resolution and approaching difficult conversations with trainees. In a workshop for women in science, the interactive exercises provided practice in conflict resolution with trainees and with senior faculty in positions of authority. In a combined workshop for faculty and staff, the exercises addressed faculty and staff issues to provide practice in defusing conflicts in the presence of a real or perceived power differential. The role-playing scenarios created for these sessions are described in the supplemental material. These single sessions were not expected to change the behavior of the participants, but served to raise awareness of approaches to manage conflict.

## Discussion

The intent of our conflict resolution training is to provide students the strategies and skillset to successfully navigate conflict. Role-playing scenarios focus on academic scenarios to prepare students for situations they may encounter in the short term with co-workers and mentors. However, these skills transcend professional boundaries as illustrated to the students by examples presented in the didactic portion of training and are useful in real life, non-professional situations. Further, this training provides a framework for managing difficult conversations around diversity, equity and inclusion.

### Reception and Effectiveness

Uniformly, across the platforms, the students felt that these sessions provided them with new information on the topic and strategies to manage conflict and negotiate. The results of these surveys suggest that these sessions are meeting the needs of graduate students. Similarly, surveys of clinicians/residents about graduate medical education leadership training demonstrate a need for instruction in the effective management of conflict and navigating difficult conversations (Fraser, Blumenthal, Bernard, & Lyasere, 2015; Hartzell, Yu, Cohee, Nelson, & Wilson, 2017). The results of studies to evaluate the effectiveness of conflict resolution training in higher education are mixed. In a graduate student conflict resolution workshop that extended over 6 sessions, the TKI and the Putnam-Wilson Organizational Communication Conflict Instrument (OCCI) were used to evaluate changes in student approaches to conflict. The pre- and post-tests were administered 10 weeks apart and the results showed a trend toward a change in approach to a cooperative style of conflict management, however the difference was not statistically significant (J. L. Brockman et al., 2010). Implementation of a conflict resolution program for graduate students and faculty in a new graduate program was evaluated by surveys and focus groups and appeared to have a positive impact upon conflict management by participants (J. Brockman, Colbert, & Hass, 2011). A two-day conflict resolution workshop for residents and medical school faculty was evaluated by surveys and a long term follow up using focus groups 12 to 18 months after the workshop (Zweibel, Goldstein, Manwaring, & Marks, 2008). The results suggest that training had an impact over the short term and importantly that elements of the training were incorporated by participants in their professional lives. Our results suggest graduate student trainees gain concepts to manage conflict from the training sessions, based upon post-test results, and that the students recognize situations where specific strategies can be applied to manage conflict months after completion of training. It is also encouraging that almost all the survey respondents indicated an intent to apply knowledge from the conflict resolution/negotiation sessions in the future.

### Workshop vs class

The first iteration of our conflict resolution sessions was delivered in a workshop format comprised of two 1.5 - 2 hour sessions with a mixture of didactic and interactive content. Incorporation of these sessions into the graduate curriculum accomplished three goals: 1) achieving broader delivery to the graduate student population, 2) delivery of the material earlier in the graduate student program and 3) extending delivery over a longer time frame. Analysis of the Michigan State University conflict resolution workshops revealed several student-identified limitations, including training a narrow cohort of students who participate in the voluntary workshops and training later, rather than earlier, in the graduate student career (J. L. Brockman et al., 2010). By extending delivery and incorporating negotiation, which shares a number of skills with conflict resolution, the exercises required retrieval practice and spaced-out practice of skills, two effective methods to promote learning and incorporation of new skills into behavior (Brown et al., 2014).

### What worked?

In both the workshop format and classroom format, students identified observing the role-playing exercises by the instructors as one of the strengths. These exercises were designed to illustrate important points or strategies and typically an example demonstrating an “incorrect” approach was illustrated by the instructors, followed by a second demonstration of a better approach using a strategy discussed in class. This method contrasted the two approaches and emphasized the differences by providing a clear example of the benefits of adopting the strategies presented in the course.

The role-playing exercises were viewed as a strength in both the workshop and traditional course format. Many students also suggested more role-playing activities would improve the course. This contrasted with a subset of students with the opposite view, that role-playing in the course should be reduced. We believe the difference reflects the comfort level of different students in role playing exercises. Likewise, in a recent survey of residents, role-playing was the least desired format in training in conflict resolution and managing difficult conversations (Fraser et al., 2015). While there are some limitations in role-playing, e.g. the interaction is artificial and it is difficult to simulate a power differential (Elliott, Kaufman, Gardner, & Burgess, 2002), most conflict resolution training utilizes role-playing (Shrader & Zaudke, 2015). Further, hands-on training and practicing skills in context are important and effective approaches in learning (Brown et al., 2014; Elliott et al., 2002).

An effective method to generate student engagement was real time polling. In the classroom, real time polling was used for brainstorming to create a list of options and this effectively translated into the virtual format via zoom in the chat feature. For example, in the negotiation portion this strategy was used to allow the students to explore the pros and cons of an Assistant Professor taking on the role of Director of Graduate studies and then preparing a list of requests they might make during negotiation with their chair. This exercise illustrated preparation for negotiation and thinking creatively in advance of negotiation.

### What didn’t work?

The role-playing exercises for conflict resolution were all designed based upon real situations experienced by our graduate students and were broadly characterized as conflicts with mentors or conflicts with peers. These engaged the students with relatable exercises for situations they may face during their graduate career. Negotiation exercises were initially more challenging. The first iteration used a negotiation scenario between teams of “students” and teams of “faculty/administrators” to negotiate a new contract between the university and the student’s union. The expectation was that the students would be engaged in discussing student issues, but this exercise was very ineffective. The second iteration used a negotiation scenario where the student played a junior faculty candidate negotiating with a departmental chair. Throughout the three sessions of the course on negotiation, students worked with each other to prepare for their negotiation. The final negotiation occurred between the students and a faculty member. Three current chairs, two ex-chairs and two senior faculty in administration played the role of chair in the simulation. This exercise also provided a unique networking opportunity for the students. A number of students identified this exercise as a strength of the course in Fall 2020.

### In-person vs Virtual Delivery

Transitioning to a virtual method of delivery during the pandemic provided some challenges. With the exception of the chat and some discussion, the format dampened the interactive nature of these sessions. Demonstrations by the instructors lacked the nuances of body language, potentially reducing the impact of illustrating different approaches to one issue. The role-playing exercises became more challenging. In the classroom, assigning roles was straightforward and instructions were provided on handouts. In the virtual format, more coordination was required and fewer instructions could effectively be delivered. In the classroom, instructors circulated around the room to monitor role playing and identifying pieces to utilize during de-briefing. The role-playing sessions in the virtual classroom were performed in breakout rooms and the movement of instructors into and out of the breakout rooms delayed progression through the class and reduced the number of interactions monitored by the instructors. In the workshop format and the Fall 2019 version of the in-class course, role playing was identified as a major strength of the course by the vast majority of the respondents to the course evaluation survey. In contrast, only one-quarter of the students in the Fall 2020 class stated that role playing was a major strength. These results suggest that the instructors’ perception of the shortcomings of role playing in the virtual classroom was shared by the students.

## Limitations

The evaluations of reaction, learning, and behavior were survey-based. The response rate for level 1 and level 2 were high (87% and 100% for the two iterations of the classroom based sessions), but the response rate for level 3 was lower (63% and 68%). While most of the respondents to the level 3 survey appeared to be aware of conflict resolution strategies and had applied or understood how to apply these strategies in a conflict they observed, the sample may not be truly representative. It is possible that trainees who had not incorporated these strategies into their management of conflict were predominant among the non-respondents and that the survey results over-represent trainees who utilized or at least recognized situations to apply these concepts. In fall 2020, when the pre-test and post-test were conducted, there was a drop in respondents to the post-test. The post-test result sample may not be truly representative of the entire cohort of trainees. There are also inherent limitations to the Kirkpatrick model, particularly in level 3 and level 4 evaluations since additional factors besides training might impact behavior and results (Holton III, 1996).

### Implementation of conflict resolution/negotiation training

The key to implementing a successful conflict resolution training program is the commitment of the faculty and administration. Expertise or previous training in conflict resolution would be an asset, but is not essential providing the faculty is willing to learn. In this case, participation in a workshop like the Train the Trainers Workshop sponsored by the Office of Intramural Training at the National Institutes of Health and consultation with experts in conflict/mediation would be useful. Professor William Rhee, a mediation expert in the WVU School of Law, facilitated our first workshop. An investment of time to research and collect material for presentation is required. This is comparable to preparation of a lecture in your general field, but outside your area of expertise. Additional time and effort are required if role-playing scenarios and other activities need to be developed. Our training sessions utilize two committed faculty members providing the opportunity for role-playing demonstrations. Additional individuals, either faculty or students who had already participated in a workshop or the class, also helped facilitate role-playing. For a group of thirty students, three or four individuals could facilitate the role-playing activities. In addition, breaking into groups of three where two students play roles and the third functions as an observer is also an effective strategy to facilitate discussion during debriefing after the role-play. The primary cost associated with implementing a conflict resolution program is faculty and staff time to develop and deliver the program.

### Future directions

During the development and implementation of this program three different formats were employed, partly by choice and partly dictated by circumstance. While we did not set out to evaluate the effectiveness of different training formats, each may fulfill different needs in future. The classroom format provides more opportunities for simulations, which is an important mechanism to reinforce concepts and adapt behavior. The workshop format may be desired by individuals with limitations on their time precluding participation in a longer course (e.g. faculty and staff). The virtual format provides an opportunity for remote instruction on conflict resolution. Our experience revealed limitations on interactions in the virtual format, but these could be overcome by adapting additional mechanisms. The chat feature provided effective interaction and additional platforms allowing real time sharing of notes or ideas might be employed. Additional strategies employing reflective exercises and/or journal activities could also be incorporated. Regardless of the format, we believe that this curriculum is valuable to trainees and are encouraged that our observations appear to indicate that training is effective.

## Acknowledgements

Professor William Rhee of the WVU School of Law was instrumental in the development and delivery of the first iteration of the conflict resolution workshop in Fall 2018. We thank William Rhee for his insightful comments during preparation of the manuscript. MS is the director of the Cell & Molecular Biology and Biomedical Engineering Training Program (T32 GM133369).

## Supporting Material

**S1 File. Professional Skills Syllabus and Conflict Resolution Scenarios**.

