## Supplemental Material for "Introducing conflict resolution and negotiation training into a biomedical sciences graduate curriculum"

##### Syllabus for SPTP: Professional Skills Development (BIOC 793A)

**BIOC 793A:**

These sessions use a combination of didactic instruction and active student participation.

---------------------------------------------------------------------------------------------------------------------

**Meeting Times:**

Weekly one hour meeting on Fridays at 3 PM in HSC-North G-119B.

---------------------------------------------------------------------------------------------------------------------

**Course Coordinators:**

Mike Schaller and Mandy Ammer

**Faculty Instructors:**

TBA for 2020

2019 instructors - David Smith, Taylor Thomas, Fatimah Matalkah, Lori Hazlehurst, Mark McLaughlin, David Klinke, Aaron Robart, Courtney DeVries

---------------------------------------------------------------------------------------------------------------------

**Course Objectives:**

The purpose of this course is to provide the opportunity to learn and practice skills that are required for the successful professional throughout their career. These skills will be valuable for professionals working in the sciences as well as other fields. Areas of instruction will include communication, conflict resolution, negotiation, working in teams and project management.

Specific Objectives

- Acquire knowledge on select topics to enhance professional development
- Develop critical professional skills
- Practice professional skills with peers
- Reflect on your developing professional skills through peer and instructor feedback

---------------------------------------------------------------------------------------------------------------------

**Course Format:**

The course contains didactic lecture components. Some didactic components will be interactive to engage students. The course also contains interactive activities allowing students to practice and develop select skills. Interactive activities will include feedback from peers and instructors.

---------------------------------------------------------------------------------------------------------------------

**Evaluation:**

Student evaluations will be based on attendance and participation in classroom activities. The grading scale is pass/fail.

---------------------------------------------------------------------------------------------------------------------

**Attendance policy:**

Students are expected to attend all sessions. If a student must miss a class, they must communicate with the instructor for that specific class PRIOR to the class to receive an excused absence. Accumulation of unexcused absences will affect the student’s final grade.

---------------------------------------------------------------------------------------------------------------------

**Prerequistes:**

Permission of the coordinator.

---------------------------------------------------------------------------------------------------------------------

**Social Justice Statement:**

The West Virginia University community is committed to creating and fostering a positive learning and working environment based on open communication, mutual respect, and inclusion. Any suggestions as to how to further such a positive and open environment in this class will be appreciated and given serious consideration.

If you are a person with a disability and anticipate needing any type of accommodation in order to participate in this class, please advise the coordinator and make appropriate arrangements with the [Office of Accessibility Services](http://diversity.wvu.edu/oas)(304-293-6700). For more information on West Virginia University’s Diversity, Equity, and Inclusion initiatives, please see http://diversity.sandbox.wvu.edu

**Conflict Resolution Scenarios**

A series of scenarios were created for role playing exercises to practice conflict resolution skills. Some scenarios were created for the conflict resolution workshops/nanocourses, others developed for practice of specific skills in the first-year graduate curriculum, and yet others for use in faculty/staff workshops. Many of the scenarios were designed for one role to resist solving the problem, providing opportunity for multiple attempts to practice conflict resolution skills. Some of the scenarios were designed to practice collaboration to solve conflict. All of these scenarios are realistic scenarios that were described by students and faculty when asked to provide examples of situations requiring management of conflict.

**A. Scenarios created for workshops/nanocourses** (students selected the scenario they wished to role play)

**Situations related to publishing and presentation.**

1. Both you and your boss want to publish your results and must agree upon a journal for submission of your manuscript. You want to pick a journal where your current data meets the requirements, your boss wants you to complete additional experiments in order to meet the requirements for a higher tier journal.  Discuss the decision with your boss.

**As mentor, you should NOT work collaboratively to solve the conflict. It is the mentor’s responsibility to decide on the journal, choosing a lesser journal could have a negative impact on you.**

**Ideas for the mentor’s argument;**

- **This is really important work and should be published in a better journal**
- **This has to be published in a higher tier journal, I need this for tenure**
- **This has to be published in a higher tier journal, you need this for your career. This will distinguish you.**
- **How could you possibly want to sell you work short and settle for a lower tier journal?**

2. You want to share your research results in posters or talks ***at a national meeting***, but your boss is discouraging you due to the potential for the project ideas getting "scooped". Discuss with your boss and try to convince him that you should present the results.

**As mentor, you should NOT work collaboratively to solve the conflict. Under no circumstances will you allow the student to go to the meeting.**

**Ideas for the mentor’s argument;**

- **You project is not complete, if you present a big lab will take the idea and scoop us**
- **I know of a number of other labs that are working on the same problem, we can’t let them know where we are**
- **We will publish in Science, and I will get tenure**
- **We will publish in Science, and you will get a great postdoc**
- **If we get scooped, we won’t be able to publish**
- **If you get scooped, you will have to start again, and you will be setback a couple of years**
- **If you worked harder, maybe the project would be done**

3. Your boss wants you to perform some experiments to help the lab complete and publish a project. Your boss has not clearly delineated a set of standards for what qualifies for manuscript authorship for efforts (academic and practical), in collaboration or individually. In the past he has not appeared to adhere to a set of standards and has re-delegated authorship status without providing opportunities for the affected individuals to address concerns. Discuss your feelings about performing these experiments and authorship with your boss.

**As mentor, you should NOT work collaboratively to solve the conflict. You have the authority to decide authorship on the paper.**

**Ideas for the mentor’s argument;**

- **I pay your salary, and technically all data produced in the lab is mine. So I get to decide authorship of my data**
- **I reserve the right to decide authorship based on the effort you put in – work harder and longer to show me you really want this**
- **Whoever in the lab contributes more figures will get authorship**

4. You have written a draft of your manuscript and your boss won’t take the time to read your manuscript and provide you feedback. Have a discussion with your boss to convince her to read your manuscript.

**As mentor, you should NOT work collaboratively to solve the conflict. You are very busy and this manuscript is very low priority at this time.**

**Ideas for the mentor’s argument;**

- **I don’t have time to do this, I am working on another student’s paper and its more important**
- **I am so busy with this grant, that I don’t have time to read your paper**
- **It is so bad, I am really having trouble getting motivated to read it**
- **I am so busy with teaching and committee work and recruiting that I can’t read your paper**
- **I really have to prepare a bunch of lectures, my boss is really pushing me to improve my teaching**
- **There is lots of time to do this, you won’t be graduating for a while, what’s your rush**

5. You have done the majority of work on a project in the lab, but you performed your experiments in a model system. Another member of the lab repeats your experiments in a clinically relevant model and shows the data to your PI. Your PI gets excited and wants to quickly write up the data and publish the findings, with the other student as first author. Discuss this decision with your PI.

**As mentor, you should NOT work collaboratively to solve the conflict. You have the authority to decide authorship on the paper and the other student provided the most critical data.**

**Ideas for the mentor’s argument;**

- **It is my responsibility to decide on authorship for all of the work that comes out of my lab**
- **I directed and funded the project, it is my prerogative to assign authorship**
- **This is really important for the other student. She really needs this first author paper**
- **This paper will get in a much better journal because of the other student’s contributions, they make this clinically relevant. She deserves first authorship for this reason**
- **Why didn’t you do the experiments to make this clinically relevant?**
- **You are early on in your career – he is in his final year and needs this to graduate**
- **You are new and still learning techniques, so I don’t want to sit on this story until you get good enough and fast enough to repeat the data in a different model**
- **You have plenty of time to get another first author paper – you don’t need this yet**
- **I pay your salary, and technically all data produced in the lab is mine. So I get to decide authorship of my data**

**Situations related to the work environment**

6. Your boss wants you to work on a project(s) that isn’t your dissertation work/won't result in a first authorship but you/your committee want to focus on your dissertation project. Discuss the situation with your boss.

**As mentor, you should NOT work collaboratively to solve the conflict. Your student MUST work on this other project since the data is critical for a grant you are writing.**

**Ideas for the mentor’s argument;**

- **This is really important, I need this data to get a grant**
- **This is really important, I need to get this grant to get tenure**
- **You are my best student and I really need your hands on this**
- **You have the expertise to do these experiments quickly and to do them right**
- **I have fully supported you and your project and now you need to support me and do these experiments**

7. Your boss has set unreasonable standards for rigor/reproducibility of your data beyond what is acceptable in the field (more cell lines, replicates, shRNAs, etc).  Discuss with your boss and try to convince her that her standards are unreasonable.

**As mentor, you should NOT work collaboratively to solve the conflict. Your student must repeat the work as directed until you are satisfied the results are correct.**

**Ideas for the mentor’s argument;**

- **Are you trying to make a fool out of me? What happens if I publish and you are wrong?**
- **This goes against what is known in the field. You must be doing it wrong**
- **I will not publish these data until those error bars are smaller**
- **If you are too lazy to add cell lines and replicates, I will have someone else take over**

8. Your boss repeatedly asks you to train new students (rotation, undergrad, medical, etc). Further, your boss asks you to train students more often than other members of the lab.  Discuss how unfair these requests are with your boss.

**As mentor, you should NOT work collaboratively to solve the conflict. This student is your best student and new students are always well trained and quickly become productive.**

**Ideas for the mentor’s argument;**

- **This is really important for the lab**
- **But you do such a great job, all your trainees have quickly contributed to the lab**
- **This is good experience for you to continue to develop your teaching skills, there is no other opportunity for you to gain this experience**
- **This is an opportunity, if you train the student and do something collaboratively, you may get another paper**

9. You and your boss disagree about the central hypothesis of your project and the best direction to more the project forward. Discuss with your boss and attempt to resolve.

**As mentor, you should NOT work collaboratively to solve the conflict. Your student MUST understand that you know the best direction for all projects in the lab and how to get them published.**

**Ideas for the mentor’s argument;**

- **You are questioning my scientific abilities?**
- **And exactly how many papers have you published?**
- **You are too young and inexperienced to understand what I am saying**
- **I have been in this game a lot longer than you. Just do what you are told if you want to stay in this lab.**

10. You feel that your PI has certain things they need to work on - managing others in the lab, engaging more frequently, being approachable.  Discuss your feelings with your boss and encourage her to improve.

**As mentor, you should NOT work collaboratively to solve the conflict. Your student doesn’t know how to manage a lab and you are not interested in listening to his advice.**

**Ideas for the mentor’s argument;**

- **I have a lot of experience managing, certainly more than you.**
- **I know what I am doing**
- **How dare you criticize my managing ability**
- **I am not the problem, you don’t respond to good managing**
- **If you were more independent this wouldn’t be a problem**
- **If you worked harder this wouldn’t be a problem**
- **I wish you were a better student**

11. You need extra help with project development but you are concerned that when you approach your boss, he/she will feel you are a burden. Discuss the need for assistance with your boss, being prepared to respond to challenges that you are burdensome.

**As mentor, you should NOT work collaboratively to solve the conflict. You believe that the student should be more independent and needs to work this out on their own.**

**Ideas for the mentor’s argument;**

- **You need to figure out how to solve your own problems**
- **No one took me by the hand and solved my problems for me**
- **This will help you learn to be independent**
- **Why can’t you do this by yourself, my other student can**
- **All you really need to do is work harder**

12. Your boss’ managing style doesn’t match what you need.  Discuss your feelings with your boss, your needs from a mentor and position solutions to the mismatch of styles.

**As mentor, you should NOT work collaboratively to solve the conflict.**

**Ideas for the mentor’s argument;**

- **How dare you tell me I am not a good mentor**
- **I have a lot of experience mentoring, certainly more than you.**
- **I know what I am doing**
- **I am not the problem, you don’t respond to good mentoring**
- **If you were more independent this wouldn’t be a problem**
- **If you worked harder this wouldn’t be a problem**
- **I wish you were a better student**

13. You are a single graduate student and another student in the lab is a parent. The other student needs to leave every day to pick up the child from daycare. The lab is working toward a grant deadline and one of the aims is based upon the other student’s project. Your PI is pushing the group to generate additional preliminary data and keeps asking you to stay late and work on experiments because the other student has to pick up the child from daycare. Discuss your feelings with your boss, trying to find a better solution to the issue.

**As mentor, you should NOT work collaboratively to solve the conflict. Your student MUST complete the data for the grant, as the other student is not available.**

**Ideas for the mentor’s argument;**

- **What is this bothering you? It is not like you have a family waiting for you at home. Who is going to miss you, your cat?**
- **You are just being petty. It is not like she is wasting time at the bar, she is taking care of her child!**
- **You are selfish. You will understand when you have children of your own how unreasonable you are being.**
- **She cannot help it that she can’t stay later. She works hard and has other responsibilities. You are lucky you don’t have that to deal with all that.**

14. You are a junior graduate student in the lab. A hierarchy, based on seniority amongst the graduate students, exists in the lab. Lab duties are not shared equitably, with the brunt of these duties falling upon the junior students. Engage with the senior graduate student, who keeps delegating tasks to you, to discuss and solve the problem.

**As mentor, you should NOT work collaboratively to solve the conflict. The lab has always been run this way. You went through this when you first started out, so it is only fair that others go through it too.**

**Ideas for the senior student’s argument;**

- **You have to earn seniority. This is the way things have always been in this lab. Don’t rock the boat.**
- **I put in my time doing all the grunt work when I first started. Now it is your turn. Why do you feel you are special?**
- **I have more responsibilities in keeping the lab running smoothly. You are new, and so you have time to do all the ancillary work.**
- **If you don’t like the rules, quit and find a new lab**

15. You have been working in a new lab for several months. Your work requires concentration since your experiments are technically challenging. You are preparing for your proposal and a fellowship application, and you read/write at your desk in the lab during incubation times in your experiments. A senior co-worker is very talkative and frequently starts trivial conversations just to socialize. This is very distracting and interferes with your experiments, reading and writing. Discuss the issue with your co-worker and try to resolve the problem.

**As senior co-worker, you should NOT work collaboratively to solve the conflict.**

**Idea’s for the senior co-worker’s argument;**

- **I think this is your problem, no one else seems to mind**
- **Just put on your headphones**
- **This is the way this lab works, if you don’t like it you should leave**
- **This is the way I like lab, and as a senior person, it goes the way I like it**
- **Socializing is very important for a team, you are not being a team player**
- **Everyone else does this too**

16. You give a public presentation and a senior colleague challenges the validity of your data and ridicules your attempt to defend your data in the question and answer period. The incident was embarrassing and demeaning to you. Have a discussion with your colleague, explaining how you feel, and try to resolve this issue.

**As senior colleague, you should NOT work collaboratively to solve the conflict. You really believe that the data is invalid.**

**Ideas for the senior colleague’s argument;**

- **I am only trying to help you**
- **I need to call you out now, so you won’t get destroyed at a national meeting**
- **Your science is so weak**
- **You are not a very good student, there are lots of problems with your experiments**
- **If I didn’t care, I wouldn’t bother criticizing you**

**Situations related to career development**

17. You are interested in a different career (non-academic) and your boss won’t allow you time to pursue other activities that would be beneficial for your career choice, e.g. travel opportunities for career-building and/or networking opportunities. Discuss with your boss and try to convince him that this will be time well spent.

**As mentor, you should NOT work collaboratively to solve the conflict. Under no circumstances will you allow the student the time required to pursue these other activities.**

**Ideas for the mentor’s argument;**

- **You need to spend more time in the lab, not less**
- **If you spend all of your time pursuing these things, you will never graduate**
- **I am training you to be a professor, if you want another career go somewhere else**
- **You don’t work hard enough and you are trying to take an easy way out**
- **This is not my responsibility, my responsibility is to train you to be a scientist**

18. You are interested in a different career (non-academic) and your boss will not have a realistic career development discussion for after graduation (not just academia and industry positions) - teaching, writing, etc. that might allow you to use and/or build upon our scientific backgrounds ---including timelines, intermediary training steps, and location. Discuss with your boss and attempt to convince him to consider training in additional areas that would benefit you.

**As mentor, you should NOT work collaboratively to solve the conflict. Under no circumstances will you allow the student the time required to pursue these other activities.**

**Ideas for the mentor’s argument;**

- **I am training you to be an academic scientist, that is what I do best**
- **I have no idea how to help, you are on your own**
- **What a waste of time. Just focus on your science, that is the be-all and end-all of graduate work**
- **This is just a distraction that will slow your progress**

**Situations related to outside activities**

19. Your boss asked you to participate in an optional event that you feel isn't a good time investment but your boss keeps bringing it up (for example, student recruiting or recurring event that doesn’t really benefit you).  Discuss your feelings with your boss and attempt to resolve the issue.

**As mentor, you should NOT work collaboratively to solve the conflict. Under no circumstances will you allow the student the time required to pursue these other activities.**

**Ideas for the mentor’s argument;**

- **These are important institutional activities and you are obligated to help**
- **You are really good in these situations and you make the institution look good**
- **It is important for you to have a presence – it reflects well on you – it reflects well on me**
- **You have to suck it up. We all have institutional duties that we have to perform**

20. You are interested in extracurricular or community events but your boss doesn't want to allow you time to pursue them (examples; teaching certificate, stats certificate, relay for life, GSO). Discuss with your boss and try to convince her to allow you to participate.

**As mentor, you should NOT work collaboratively to solve the conflict. Under no circumstances will you allow the student the time required to pursue these other activities.**

**Ideas for the mentor’s argument;**

- **I pay you to work in the lab not to pursue other activities**
- **Time away from lab does nothing but slow down your progress**
- **This would set back completion of your degree**
- **Your project is competitive and time out of lab will get us scooped**
- **I really need you in the lab to generate the preliminary data for my grant**
- **This is really a waste of your time and will not really help advance your career**

21. Your boss does not want you to make time for out of science life priorities that you consider important for your physical and mental health. Discuss with your boss and try to convince him that these are important issues requiring resolution.

**As mentor, you should NOT work collaboratively to solve the conflict. Under no circumstances will you allow the student to take any more time off.**

**Ideas for the mentor’s argument;**

- **You don’t work hard enough and you don’t get enough done**
- **All you want to do is extracurricular, you don’t want to work**
- **You keep taking so much time off, you will be a student forever**
- **I am paying you to work in the lab, not to keep taking vacations**
- **This project is going to fail because of your work habits**
- **You are selfish and this will have a negative impact on me and others in the lab**

**Situations related to completing the degree**

22. You have completed your requirements for degree, but your boss wants you to stay in the lab and generate more data before writing up your thesis and defending your dissertation. Discuss with your boss attempting to persuade her that you are ready to write.

**As mentor, you should NOT work collaboratively to solve the conflict. Under no circumstances will you allow the student to graduate until they lay the groundwork of a project for the next student.**

**Ideas for the mentor’s argument;**

- **You are not done yet, you are done when I say you’re done**
- **Right now you are average, publish another paper and you will stand out**
- **I supported you all through your training, now you owe me this data for my next grant**
- **This project needs to be developed so the next student has something to start with**
- **I finally have you trained to be productive, now you need to produce something for me**
- **You are my best set of hands and I need you to stay for a while**

**B. Scenarios created/modified for the first year curriculum** (all students performed the same role plays with guidance to practice specific skills)

1. **“ I don’t wanna do the dishes!”**

STUDENT SCENARIO

You are a junior graduate student in the lab. A hierarchy, based on seniority amongst the graduate students, exists in the lab. Lab duties are not shared equitably, with the brunt of these duties falling upon the junior students. These duties include dishwashing and making up lab stocks of reagents. Engage with the senior graduate student, who keeps delegating you tasks, to discuss and solve the problem.

The goal of the exercise is to practice:

- Situation, action, impact feedback
- Listening
- Separating positions from interests

One person will play the junior student and the other person will play the senior student. The person playing the senior student should monitor the discussion for effective use of the skills we discussed today. We will debrief and ask for volunteers to report back effective approaches by the “junior students”.

**SENIOR STUDENT SCENARIO**

**As the senior student, you should NOT work collaboratively to solve the conflict. Under no circumstances will you allow the junior student to reorganize the duties in the lab.**

**Ideas for the senior student’s argument;**

- **The lab has always been run this way. You went through this when you first started out, so it is only fair that others go through it too.**
- **You have to earn seniority. This is the way things have always been in this lab. Don’t rock the boat.**
- **I put in my time doing all the grunt work when I first started. Now it is your turn. Why do you feel you are special?**
- **I have more responsibilities in keeping the lab running smoothly. You are new, and so you have time to do all the ancillary work.**
- **I am a senior graduate student and my work is more important than yours**
- **I won’t always be here to pick up your slack, so you need to learn how to do this before I’m gone**
- **If you don’t like the rules, quit and find a new lab**

2. **“I want to go to a scientific meeting”**

STUDENT SCENARIO

There is a national meeting in your field and your scientific hero is giving the keynote address. You want to attend the meeting, share your research results in a poster or talk, and maybe meet your hero. Your boss is opposed to you attending the meeting. Discuss this with your boss and express your strong desire to attend the meeting.

The goal of the exercise is to practice:

- Situation, action, impact feedback
- Listening
- Separating positions from interests

One person will play the student and the other person will play the mentor. The person playing the mentor should monitor the discussion for effective use of the skills we discussed today. We will debrief and ask for volunteers to report back effective approaches by the “students”.

**MENTOR SCENARIO**

**As mentor, you should NOT work collaboratively to solve the conflict. Under no circumstances will you allow the student to go to the meeting.**

**Ideas for the mentor’s Argument;**

- **You project is not complete, if you present a big lab will take the idea and scoop us**
- **I know of a number of other labs that are working on the same problem, we can’t let them know where we are**
- **We will publish in Science, and I will get tenure**
- **We will publish in Science, and you will get a great postdoc**
- **If we get scooped, we won’t be able to publish**
- **If you get scooped, you will have to start again, and you will be setback a couple of years**
- **If you worked harder, maybe the project would be done**

3. **“Where’s my blot?”**

**Side A**: Upon walking in to the lab on a sunny morning, you notice that your lab partner left a mess. There are antibodies out on the counter, a pile of dishes in the sink, and they forgot to tear down your protein gel and set up the transfer apparatus like they promised they would. Two days of work is ruined, and the antibodies you needed for your blot are also ruined. This is not the first time your lab partner has let you down, and now you have nothing to show your boss for your weekly lab meeting.

In bullets below, sketch out your “1^st^ story”

**Side B**: The previous night, you were working in the lab, waiting on your lab partner’s gel to finish so you can set up the transfer. You get a call from your baby sitter, and your child is sick, vomiting, and running a high fever. You dash out of the lab, forgetting everything to get there so you can take your child to Urgent Care. You got to your house around midnight, and by the time you got your child settled, you don’t get to sleep until after 3AM. You realize that you let your lab partner down, but some things are more important than a blot.

In bullets below, sketch out your “1^st^ story”

Side A student explains their 1^st^ story, then Side B student explains their story.

Together write out the 3^rd^ story (objective story from a 3^rd^ party’s perspective)

Interesting follow up point:

Person 2 is a woman in my scenario – Does your view of what happened with the sick kid change if I tell you it is a man?

4. **A difficult conversation – Engaging a reluctant participant in discussion**

You are an Assistant Professor. Your R01 was reviewed and discussed at study section, but was not funded. The reviewers criticized the feasibility of your project and indicated significant additional preliminary data is required. Your deadline for resubmission is approaching. You have been pushing your graduate student to generate the data required.

Your graduate student comes to talk with you and informs you that they are quitting graduate school. They are very reluctant to talk.

Have a discussion with your graduate student using the following techniques to engage:

“me-me, and”

Contrasting

Tentative statements

Asking

Mirroring

Paraphrasing

**Graduate student – very reluctantly engage in conversation.**

**C. Scenarios created for faculty/staff workshops**

**Situations with trainees**

1. The three students in your lab have done all of the work on a project that you are getting ready to publish. One of the students laid all of the ground work and made all of the cell lines/mutants which were used extensively throughout the paper, while the other students ran experiments that were used in the figures of the paper. Each student feels that they should be first author, and all of them are against a triple-first author caveat. Things are getting nasty in the lab and no one is talking to anyone else. Discuss this situation with your students.

**As student in the lab, you should NOT work collaboratively to solve the conflict.**

**Ideas for Student’s argument:**

- **I am senior in the laboratory, and I need to graduate before the others. I need this to graduate; therefore, I should be first author.**
- **Without my cloning expertise, this project would have failed from the start.**
- **I put in so many hours on this, and if I don’t get a first authorship, I think I may quit.**
- **You are just playing favorites. I know you don’t like me as much as the others in the lab, and I should know how you are by now and not continually be disappointment in your treatment.**

2. You have a student who has been in the lab for over 2 years. Recently, you have noticed the student acting a little off, and their appearance and overall attitude has changed and the student is becoming more withdrawn. You find that you are having difficulty communicating with the student, and all you do is argue and fight over the smallest things during your weekly lab meetings. You realize that the lab has been crazy trying to get data to finalize your figures for your grant. Finally, the student approaches you and tells you that they are miserable and would like to quit your lab. This would be catastrophic to you and your progress. Discuss the situation with your student.

**As student in the lab, you should NOT work collaboratively to solve the conflict.**

**Ideas for Student’s argument:**

- **You only care about publishing papers and about your career – you are killing my will to be a scientist.**
- **You don’t understand how hard this work is – you stay in your office and read all day.**
- **All you care about is yourself and your career. I am nothing more than a hired hand to you.**
- **I don’t think I want to spend the rest of my life feeling this stressed and hurried all the time.**
- **I have never had a problem with anxiety – now I am on medication for it. You are ruining my life.**

3. You recently hired a post-doc that came highly recommended by one of your colleagues. While this post-doc is proficient and very knowledgeable, you feel that they are not working as hard in the lab as they should be. They continually take long lunches, and they often show up at 11AM to start their day. While they claim they get most of their work done late at night, you can see they are not finishing experiments in a timely fashion, and they are missing official deadlines. How do you discuss this issue with your post-doc?

**As the post-doc, you should NOT work collaboratively to solve the conflict.**

**Ideas for Post-doc argument:**

- **I am working as hard as I can. You expect me to help the graduate students when they are having issues. Make up your mind – do I only do my work or do I help out in the lab.**
- **I am doing a lot of trouble-shooting. I don’t bring you all of my negative data.**
- **You are never here past 6pm. How do you know if I am working hard all night or not?**
- **I know I missed a deadline, but you can always fix that for me – you are friends with the chair, right?**
- **I am trying – I have a young family and I would like to have a life.**
- **Science is harder now than when you were a post-doc. Why are you not more supportive?**

**Situations with authority figures**

4. **“New uncompensated responsibilities – faculty version”** (resistance to solving the problem)

You are a newly appointed Assistant Professor in your department. You recently wrote a small grant and it was funded. You would like to expand your research lab, and this grant will help you to attract graduate students and to be able to hire a technician. However, you have a large teaching load and you also have your service duties. You would like to approach your chair to discuss the possibility of getting some release time, or having some of your time protected for your research. Discuss your wants with your chair.

**As the Chair, you should NOT work collaboratively to solve the conflict.**

**Ideas for Chair’s argument:**

- **You signed a contract when you were hired that you would teach a certain number of classes.**
- **Congratulations on your grant. If it was bigger, I would consider your request. Go and get a bigger grant and we can talk.**
- **All of your colleagues are in the same boat, and they manage their time just fine. Maybe you need to take a time management class.**
- **What do you suggest? That I teach your classes?**

5. **“New uncompensated responsibilities – faculty version”** (collaboration in conflict resolution)

**Role 1:** You are an Assistant Professor. You were protected for six months to develop your research program. For the last 18 months you have been teaching a full load in the undergraduate and graduate curriculum in the department. There are three graduate students in your lab, you have one senior author paper and your NSF grant was just funded. Your boss wants to talk with you about becoming the Director of Graduate Studies for your graduate program (this is no additional compensation for this added responsibility).

Engage in the conversation with the Chair of the department about this additional responsibility.

**Role 2:** You are the Chair of the department. One of your successful faculty members has just left to take a position at another institution. This faculty was the Director of Graduate Studies for the program and you need to quickly identify a replacement for this position as the new students will be entering the program in a few months. There are several senior faculty members of the department, who are not interested in the position and any commitment would only be short term. You have other faculty whose personnel management skills are not suitable for the position. Several other faculty are struggling academically and you are concerned about their future. The best candidate is an Assistant Professor who has been at the institution for two years, is effectively mentoring 3 graduate students in the lab, has published and was just awarded an NSF grant.

Engage in the conversation with the Assistant Professor about taking on this responsibility. Unfortunately, your budget will not allow additional compensation for the Assistant Professor.

6. **“New uncompensated responsibilities – staff version”**

**Role 1:** You are the newest member of an administrative team and have been working as a member of the team for the past year. You rapidly acclimatized to your position, established good relationships throughout the department and have mastered all of your duties. Further, you have modified the procedures associated with your duties to significantly increase the efficiency of the performance of your duties. Your boss wants to talk to you about assuming a bigger role in the unit and the first task, which will be reviewing and modifying all procedures in the unit to increase efficiency. This will require you to dictate to others how they should do their jobs (there is no additional compensation for this added responsibility).

Engage in the discussion with your boss to express your thoughts and concerns about this larger role in the unit.

**Role 2:** You are the head of the unit and the newest member of your administrative team has been performing exceptionally. The team member has established good relationships within the department and taken the initiative to modify procedures affiliated with his/her duties to significantly enhance efficiency. You have been concerned about outdated procedures in the department, the impact upon efficiency of operation and impact upon morale. You would like to task the newest team member to review and modify all departmental procedures.

Engage in the conversation with the team member to discuss your plans and alleviate concerns. Unfortunately, your budget will not allow an increase in compensation for the increased responsibility.

**Situations with colleagues**

7. **“Collaboration…….or not”**

**Role 1**: You are an Assistant Professor. You were protected for six months to develop your research program. In the last 12 months you have begun teaching in the both the undergraduate and graduate curriculum in your department. You have one graduate student in the lab and you are not yet funded. You have been discussing your main project with a senior faculty member, who is interested in collaborating with you. At a faculty gathering to brain storm about grants, your senior colleague presents “his/her ideas”, which includes key experiments you plan as part of your main project (no collaboration is mentioned).

Approach your senior colleague to discuss the issue. Try to use the situation impact feedback and active listening techniques to express yourself and learn more about your colleague’s motivation.

**Role 2**: You are a well-established senior Professor. For many years you have been considering an important problem, but only intellectually, and you have not initiated a project to tackle the problem. An Assistant Professor has been on campus for a couple of years and has a strong interest in a related problem. You have had many conversations with the Asst Prof, which have been very stimulating. You have contributed important ideas to him/her and the conversations have driven an evolution in your thinking about the problem. You are very excited about these ideas and have begun planning a project. At a recent faculty gathering to brain storm about grants, you outline your ideas and present thoughts on key experiments in your proposed project.

The Assistant Professor appears upset after your presentation and approaches you to discuss a problem. Try to use active listening techniques to learn more about your colleague’s perceptions of what happened.

8. **“Holistic or merit-based approaches”**

**Role 1:** You are on a committee that will establish new school-wide hiring policies and guidelines. You believe that diversity is an asset, that diverse communities provide more creative environments and that increasing diversity is critical for the long-term success of the college. You support a holistic approach to hiring, where life experiences are factored into decisions, in addition to academic credentials and past success. There are two groups on your committee, one supportive of the holistic approach and the other who believe that credentials and past success best predict future success and only by hiring the best candidate under these criteria will the school succeed in the long term.

Engage a colleague from the other camp to discuss your differing views on hiring. Try to use acknowledgement, getting past positions to interests, reframing arguments, me-me and.

**Role 2:** You are on a committee that will establish new school-wide hiring policies and guidelines. You believe the reputation of the School is static or declining and that re-invigoration through hiring is critical to correct this trend. This will require hiring the brightest, hardest working and most likely to succeed people. You believe that hiring should be merit based and that academic measurements, e.g. papers, fellowships and grants, provide the best measurements of merit and are the strongest indicators of future success. Some colleagues on the committee share your views while others believe a holistic approach, where life experiences are factored into decisions, in addition to academic credentials and past success, will be better for the institution in the long run.

Engage a colleague who supports holistic admissions to discuss your differing views on hiring. Try to use acknowledgement, getting past positions to interests, reframing arguments, me-me and.

9. **“You are taking advantage of me!”**

**Role 1:** You are a new member of an administrative team, but have extensive administrative experience from past positions. Another member of the team has been part of the administration for many years and is well respected. The two of you have been partnered to review the entire operation of the unit, evaluate strengths and areas requiring improvement, develop a plan to modify operations to increase efficiency and a budget to implement the changes. Your colleague has taken the lead to begin drafting recommendations without consulting you and has had two informal meetings with the boss to discuss progress. You are concerned that your colleague is taking advantage of what was supposed to be an equal partnership.

Approach your senior colleague to discuss the issue. Try to use the situation impact feedback and active listening techniques to express yourself and learn more about your colleague’s motivation.

**Role 2:** You are the senior member of an administrative team and have many years’ experience and well-established relationships in the unit. A new member, who has extensive administrative experience from past positions, has joined the team. The two of you have been partnered to review the entire operation of the unit, evaluate strengths and areas requiring improvement, develop a plan to modify operations to increase efficiency and a budget to implement the changes. It is a little unusual for a new team member to be assigned such a large role before establishing themselves as part of the team. Since you have more familiarity with the unit, you independently begin drafting recommendations that you consider “no-brainers”. As you have good relationships in the unit, including with the boss, you have had informal discussions about your ideas with the boss.

Your junior colleague asks about the equality of the partnerships. Engage in discussion with your colleague. Try to use active listening techniques to engage your colleague and use steps for effective communication.
